# Chiral vortex dynamics on membranes is an intrinsic property of FtsZ, driven by GTP hydrolysis

**DOI:** 10.1101/079533

**Authors:** Diego Ramirez, Daniela A. García-Soriano, Ana Raso, Mario Feingold, Germán Rivas, Petra Schwille

**Affiliations:** Department of Cellular and Molecular Biophysics, Max Planck Institute for Biochemistry, Martinsried, Germany; Graduate School for Quantitative Biosciences (QBM), Ludwig-Maximillians-University, Munich, Germany; Centro de Investigaciones Biológicas, Consejo Superior de Investigaciones Científicas (CSIC), Madrid, Spain; Department of Physics, Ben Gurion University, Beer Sheva, Israel

**Keywords:** Self-organization, cytomotive filaments, Z-ring, bacterial division, TIRFM

## Abstract

The primary protein of the bacterial Z ring guiding cell division, FtsZ, has recently been shown to engage in intriguing self-organization together with one of its natural membrane anchors, FtsA. When co-reconstituted on flat supported membranes, these proteins assemble into dynamic chiral vortices whose diameters resemble the cell circumference. These dynamics are due to treadmilling polar FtsZ filaments, supposedly destabilized by the co-polymerizing membrane adaptor FtsA, thus catalysing their turnover. Here we show that FtsA is in fact dispensable and that the phenomenon is an intrinsic property of FtsZ alone when supplemented with a membrane anchor. The emergence of these chiral dynamic patterns is critically dependent on GTP concentration and FtsZ surface densities, in agreement with theoretical predictions. The interplay of membrane tethering, GTP binding, and hydrolysis promotes both, the assembly and the destabilization of FtsZ polymers, leading to the observed treadmilling dynamics. Notably, the vortex chirality is defined by the position of the membrane targeting sequence (mts) and can be inverted when attaching it to the opposite end of FtsZ. This reveals a so far unknown vectorial character of these cytomotive filaments, comprising three orthogonal directions: Filament polarity, curvature, and membrane attachment.

## Introduction

The FtsZ protein, a self-assembling GTPase present in the cytoplasm of most bacteria, is a tubulin homologue and the main component of the *Escherichia coli* divisome, the molecular machinery driving cytokinesis^1,2^. It interacts with additional proteins forming a dynamic ring at the cytoplasmic membrane, which acts as a scaffold to recruit the remaining elements of the divisome. *In vitro* studies using the FtsZ protein from *E. coli* have shown that GTP drives the (Mg^2+^-linked) concerted assembly of FtsZ monomers to form narrow-size distributed single stranded filaments that are flexible and tend to further associate into higher order polymers sufficiently plastic to adopt multiple geometries (multi-stranded fibers, bundles, circles, toroids, etc) depending upon experimental conditions^3,4^. Like other cytoskeletal filaments, these are dynamic with a GTP-dependent turnover of FtsZ, which is considered to be central for positioning and function of the Z ring^3,5,6^.

For membrane attachment, FtsZ requires two additional divisome proteins, FtsA and ZipA. Together they form the so-called proto-ring, the first molecular assembly of the divisome^7,2^. FtsA is an amphitropic protein that associates to the membrane by an ATP-linked process mediated by a short amphipathic helix^8^. The bitopic membrane protein ZipA contains a short N-terminal intracellular region, a trans-membrane region, and an extracellular C-terminal FtsZ-interacting domain connected by a flexible linker region^9^. FtsZ binds the proto-ring tethering elements through its C-terminal end, which it is also the interaction region for FtsZ-regulating proteins such as MinC, inhibiting FtsZ polymerization and hence FtsZ ring formation at undesired locations. Thus, the C-terminal region of FtsZ acts as a central hub integrating signals that modulate divisome assembly in *E. coli*^2^.

Minimal membrane systems, from nanodiscs and supported bilayers to free standing vesicles, have been used as scaffolds to reconstitute different combinations of proto-ring subsets in order to test their functional properties^10,11^. In this way, an FtsZ chimera, FtsZ-YFP-mts, where the FtsZ central hub was replaced by YFP and an amphipathic helix to provide autonomous membrane attachment, was found to become internalized and accumulated in narrow regions of tubular liposomes, forming static ring-like structures. The polymerization of the chimeric FtsZ protein at the external face of liposomes induced their deformation and tubulation^12,13^. FtsZ static assemblies were also found when wild type FtsZ and FtsA were co-reconstituted inside liposomes^14^. Furthermore, wtFtsZ assembly inside ZipA-containing permeable giant vesicles yielded limited dynamic structures resulting in vesicle shrinkage^15^.

Reconstitution on supported bilayers has allowed studying the structural organization and dynamics of proto-ring elements at the membrane using surface-sensitive techniques^16, 11^. They have been used to show that membrane-tethered FtsZ, either by mts or an extracellular variant of ZipA, assembles at the bilayer forming dynamic polymer networks that can reorganize by fragmentation, annealing and lateral condensation^17,18^. The coupling between FtsZ and the site-selection MinCDE complex has also been investigated in supported and micro-engineered bilayers^6,19^. There it has also been shown that the spatial regulation of FtsZ by MinC is crucially dependent on the turnover of FtsZ within the filaments.

Recently, the co-reconstitution of wtFtsZ and FtsA on supported bilayers has revealed that FtsA promotes the self-organization of FtsZ fibers into dynamic patterns, giving rise to coordinated streams and swirling rings with preferential directions, due to treadmilling dynamics, as revealed by total internal reflection fluorescence microscopy, TIRFM^17^. In contrast, this dynamic behaviour was not observed when FtsZ was tethered to the membrane through ZipA, or when the membrane targeted FtsZ variant was used. These observations suggest that FtsA plays a crucial role in these dynamics, not only recruiting FtsZ polymers to the membrane but also destabilizing them by biochemical mechanisms yet to be identified.

In order to elucidate these treadmilling dynamics more quantitatively, and to particularly investigate their dependence on chemical energy provided by nucleotide hydrolysis, we set out to study FtsZ vortices in a minimalistic setting. Strikingly, we found that in contrast to what has been reported previously, FtsA, and along with it, ATP, are in fact dispensable components. Under certain conditions as outlined below, membrane-targeting of FtsZ is sufficient for the protein to unfold its spatial self-organization behavior, resulting in swirling rings on flat supported membranes. GTP was found to promote the dynamic association/dissociation of FtsZ polymers, which increasingly takes the shapes of treadmilling rings, rotating with a GTP-dependent velocity of up to 4.78°sec^−1^. Remarkably, the ring diameter was conserved at ca. 1µm as reported earlier, however we report a time-dependent ripening and thickening of the individual rings. To reveal the origin of the distinct chirality of the vortices, we used an FtsZ mutant version with the mts cloned to the opposite end of the protein and could thereby revert the direction of rotation. This indicates that membrane attachment constitutes an orthogonal vector to both, the filament growth direction, and the direction of curvature. Together with the apparent thickening of the rings, this could point towards a possible physiological role of this striking self-organization phenomenon.

## Results

The protein chimera FtsZ-YFP-mts (0.5 µM) in its GDP-bound form (corresponding to a non-assembled state, according to sedimentation velocity, **Fig. S1**) did not form visible structures on a supported lipid membrane, as revealed by TIRFM. However, the protein in its monomeric form is either attached to or in close proximity to the membrane, as observed by the high intensity on the membrane (**Fig. S2**). Addition of GTP (4 mM) and magnesium (5 mM) to promote protein polymerization (**Fig. S1**) resulted in the appearance of FtsZ filaments at the membrane, followed by self-organization to form dynamic ring-like structures (**Fig. 1C, Movie S1**). Right after adding GTP, the bulk intensity on the membrane decreased (**Fig. 1B,C**). In the first 5 minutes, highly dynamic short filaments were observed, which showed lateral diffusion, bending and straightening (**Fig. 1B,C**). After 10 min, those short filaments formed small and dim closed circular structures that tended to be highly unstable; closed filaments were able to open, diffuse and fuse with adjacent filaments or to close back. At later times, closed circular strands turned into ring-like structures with a defined center (**Fig. 1B,C**). Every ring was apparently connected to adjacent rings by highly motile strands that created a network of connecting rings.

**FIGURE 1.**
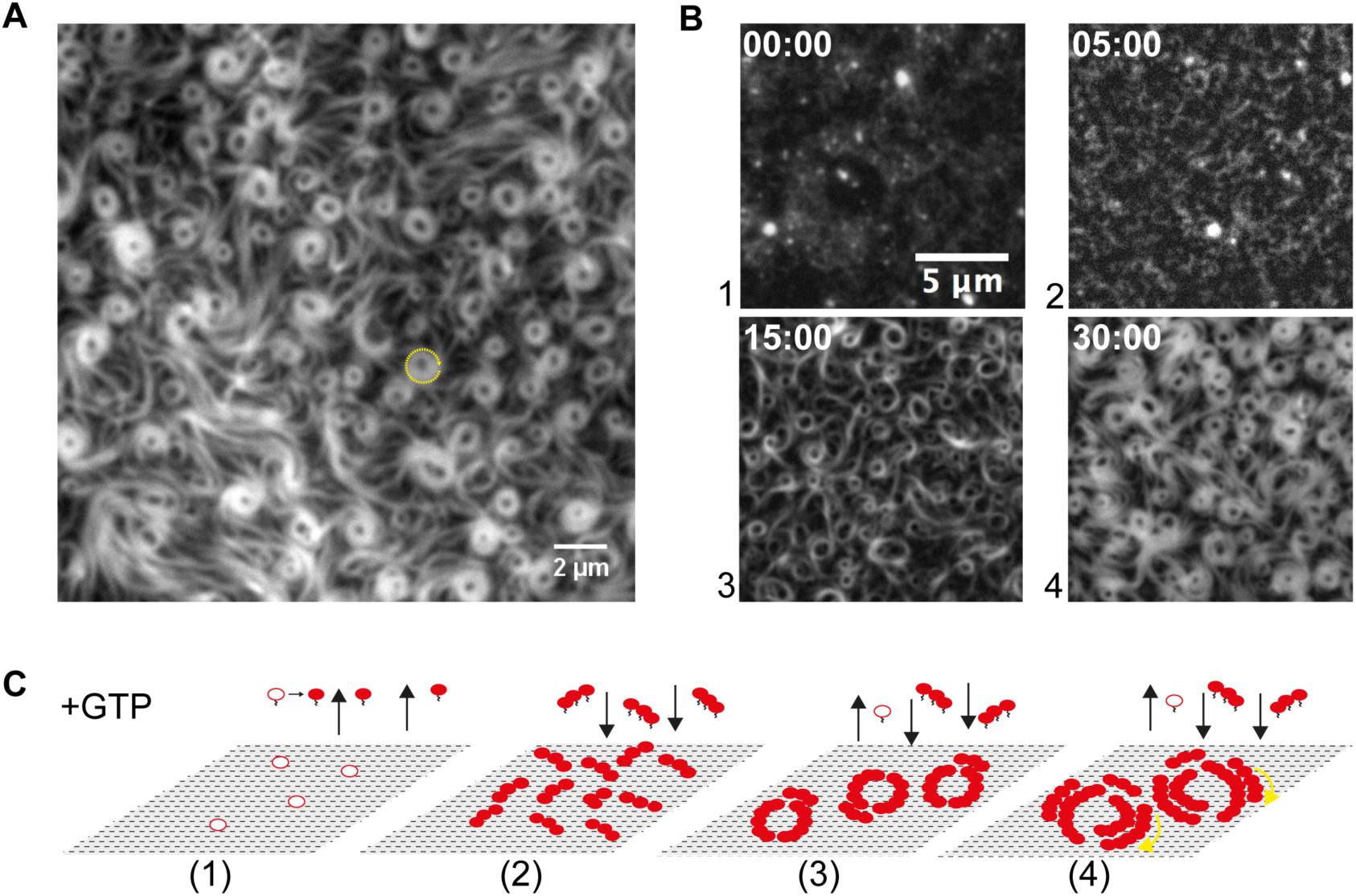
Membrane-targeted FtsZ is able to form dynamic cytoskeletal patterns on supported lipid bilayers. (A) FtsZ-YFP-mts assembles into circular vortices, which consistently rotate clockwise. (B) Representative snapshots from a time-lapse experiment displaying different stages of ring formation. Images were taken every 10 seconds using TIRF illumination (YFP channel). After GTP addition, monomers are released into solution where the polymerization is triggered (0:00 min). Then, short protofilaments from solution slowly return to the membrane forming small structures with high lateral diffusion (5 min) (**Movie S1**). After 10 min, those short filaments formed unstable and small circular structures that transiently open and close. At later times, ring-like structures with a defined center become predominant (15 min). In the long time regime (30 min), rings become larger and thicker. (C) Schematic representation of the main events described in (B). White spheres represent FtsZ-YFP-mts in its GDP state. Red spheres correspond to FtsZ-YFP-mts in the GTP state.

Single rings appeared as rotating vortices, meaning that the light intensity along the perimeter of the ring changed periodically in time. In addition, these vortices consistently showed a chiral clockwise rotation (**Fig. 2A**). The directional ring dynamics was confirmed by the positive slope tendency of kymographs generated along the ring circumference, which allowed estimating the rotational speed of these structures to be around 2.5 µm/min or 4.8°sec^−1^ implying that one full rotation takes 1.3 min **(Fig. 2A)**. To investigate whether the emergence of the collective streams and swirls was an intrinsic property of the membrane-targeted FtsZ dynamics linked to GTP hydrolysis, we carried out similar self-organization assays using a variant of the FtsZ chimera with no GTPase activity (**Fig. S3**), in which the Threonine at position 108 was replaced by an Alanine (FtsZ*[T108A]-YFP-mts). At a concentration of 0.2 µM and 30 min after the addition of GTP and magnesium, well-defined rings similar in size to the ones found with FtsZ-YFP-mts could be observed (**Fig. 2B**). Interestingly, these rings did not seem to rotate (**Movie S2**), a feature supported by the lack of clear patterns in the kymographs generated to track polymer dynamics (**Fig. 2B**). From these results we concluded that the GTPase activity of FtsZ is involved in the formation of the dynamic patterns.

**FIGURE 2.**
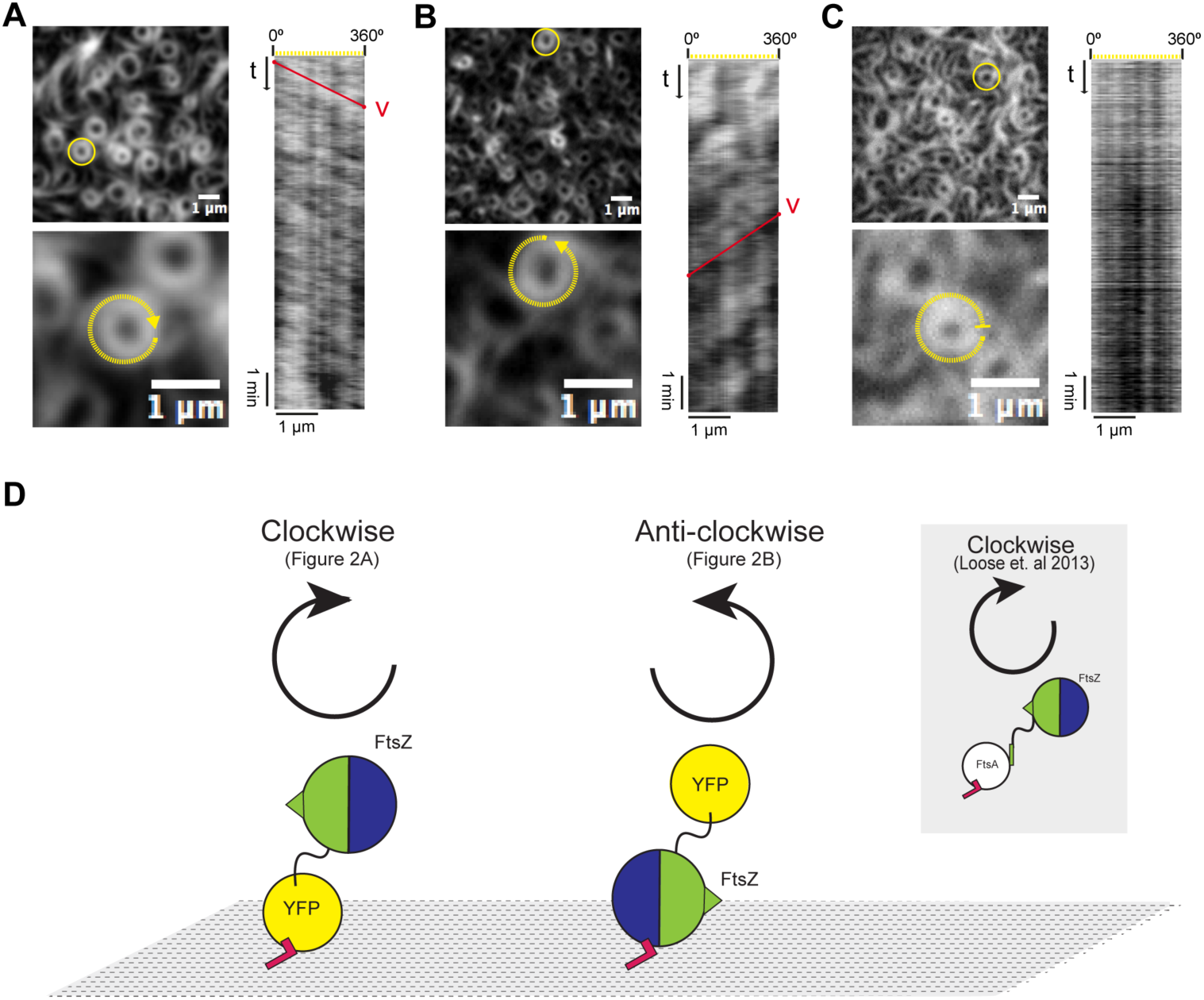
Dynamic behavior and chirality of FtsZ rotating vortices depend on GTPase activity and location of the membrane targeting sequence (mts). (A) Representative snapshot of FtsZ-YFP-mts (mts in the C-terminus) at 4 mM GTP. On the right, kymograph analysis shows a positive slope that corresponds to the apparent clockwise rotation time of the selected ring (yellow circle). (B) Representative snapshot of mts-H- FtsZ-YFP (mts in the N-terminus) at 4 mM GTP. The kymograph analysis shows a negative slope that corresponds to an apparent anti-clockwise rotation time. This indicates that the position of the membrane targeting sequence determines the chirality of the apparent rotation. (C) FtsZ*[T108A]-YFP-mts rings. The kymograph analysis shows no apparent slope in contrast with the kymographs showed in (A) and (B). The absence of GTPase activity generates static rings, suggesting that the apparent rotation in (A) and (B) is mediated by GTP hydrolysis. (D) FtsZ is tethered to the membrane through FtsA (right diagram); the C-terminal peptide of FtsZ (central hub) binds FtsA, and the latter anchors the membrane via its amphipathic helix (red polygon). The C and N domains of FtsZ are represented as blue and green hemispheres, respectively. In the case of FtsZ-YFP-mts, the FtsZ binding peptide is replaced with YFP and an amphipathic helix (left diagram). In mts-H-FtsZ-YFP, this amphipathic segment is incorporated at the N-terminal of FtsZ, while the YFP remains at the C-terminal (mid diagram).

Next, we reasoned how the obvious chirality, resulting in only clockwise rotation of the rings, could be established. Since it would have been too circumstantial to fully revert the polarity of the protein, we simply reverted the membrane attachment to the opposite, N-terminal, end (mts-H-FtsZ-YFP). Again, at a concentration of 1.25 µM and after 10 minutes of GTP and magnesium addition, defined and dynamic rings were observed. But now, strikingly, the swirls appeared to rotate in the anti-clockwise direction (**Movie S3**). The anti-clockwise directionality was confirmed by the negative slope tendency of the kymographs. The rotational speed of these patterns was estimated to be around 1.4 µm/min or 2.7°sec^−1^ implying that one full rotation takes about 2.2 minutes (**Fig. 2C**). From these observations we conclude that the mts positioning and thus, the membrane attachment domain, determines the direction of apparent rotation and thus, the direction of polymerization in the membrane plane.

To validate the concentration dependence of vortex formation, we studied the structures formed by FtsZ-YFP-mts on the membrane at a fixed concentration of GTP (4mM) and different protein concentrations (between 0.1 and 1.0 µM). At 0.1 µM, no polymers could be detected (**Fig. S4**), while at the highest protein concentration tested (1.0 µM) very abundant three-dimensional polymer networks at the vicinity of the membrane, but no dynamic rings, were observed (data not shown). We found that 0.2 µM was the minimal protein concentration at which dynamic structures on the bilayers could be visualized (**Fig. 2A**). Next, we compared the kinetics of protein adsorption at 0.2 µM with the one measured at 0.5 µM (**Fig. 3A**), conditions previously used to detect the swirling rings (**see** **Fig. 1**). Immediately after addition of GTP, a similar membrane adsorption rate was found at the two protein concentrations (**Fig. 3A**); short and highly dynamic filaments were evidenced initially (**Fig. 3B**). The transition from short filaments to rudimentary circular structures (grey area in **Fig. 3A**) occurred at similar times for the two protein concentrations assayed. After 10 min, the adsorption rate slows down at 0.2 µM, compared to 0.5 µM where it still increases, suggesting that the kinetics of ring stabilization and widening of the structures was concentration limited (**Fig. 3A**). The difference between the behaviours at 0.2 µM and 0.5 µM FtsZ-YFP-mts for t > 10 min is further emphasized by the corresponding dynamics of the protein pattern morphology (**Fig. 3B**). In particular, the ring morphology observed at t = 45 min for 0.2 µM protein (**Fig. 3B**, lower right panel) is qualitatively equivalent to that at t = 20 min for 0.5 µM (**Fig. 3B**, upper mid panel). This morphological similarity correlates with the equal surface protein concentration at these two time points (points 2 and 3 in **Fig. 3A**), showing that this is the factor determining the nature of the protein network on the membrane. To further establish the correspondence between the morphologies at time points 2 and 3 of **Fig. 3A**, we analysed the distribution the corresponding ring diameters. These distributions were found to be practically identical with average diameters of 0.98 +/- 0.14 µm (0.2µM) and 0.94 +/- 0.16 µm (0.5 µM) (**Fig. 3B**), suggesting that although the adsorption rates were different at these times, the proteins condensed into similar structures.

**FIGURE 3.**
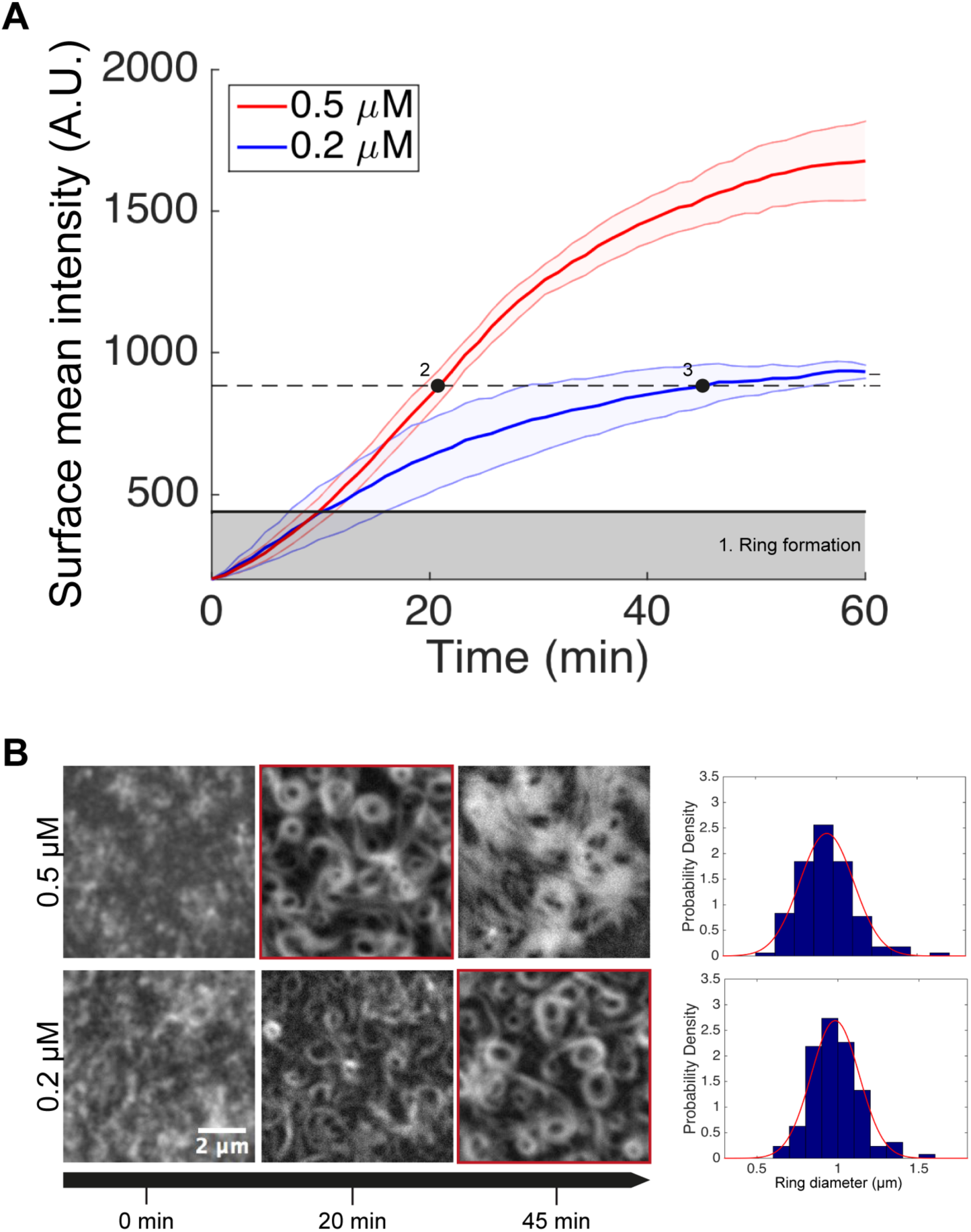
FtsZ-YFP-mts ring formation depends on protein surface concentration. (A) Time-dependence of the average FtsZ-YFP-mts fluorescence from the membrane at two different protein concentrations (fluorescence intensity is proportional to the amount of protein on the bilayer). Initially, in the range of intensities below 450 A.U> (grey area) short protein filaments diffuse laterally forming unstable rings that open and close. Later, stable rings with well-defined center and diameter start assembling, such that at intensities around 880 A.U. (dashed line) the filaments form a network of chiral vortices. While for a protein concentration of 0.2 µM, the system reaches this regime at t = 45 min (time point 3), at 0.5 µM it only takes ~20 min (time point 2). (B) Images from a time-lapse experiment corresponding to t = 0, 20, 45 min for each of the two protein concentrations, 0.2 µM and 0.5 µM, together with the corresponding ring diameter distributions at time points 3 and 2, respectively (see (A)). Note that, at t = 20 min, the protein network corresponding to 0.2 µM FtsZ-YFP-mts (lower mid panel, only few rings together with thin filaments) has a different appearance than that at 0.5 µM (upper mid panel, red border, many thick rings). In contrast, the protein pattern at t = 45 min and 0.2 µM FtsZ-YFP-mts (lower right panel) is similar to that at t = 20 min and 0.5 µM FtsZ-YFP-mts (upper mid panel, red border), showing that the main determinant of the nature of the network is the protein density on the bilayer.

As the levels of GTP modulate the assembly and disassembly of FtsZ polymers mediated by GTP hydrolysis, we then followed the formation of the dynamic patterns at a fixed protein concentration (0.2 µM) using increasing GTP from 0.04 to 4 mM. In contrast to the dynamic vortices found at 4 mM GTP (**Fig. 1**), highly ordered filaments were rapidly formed at 0.04 and 0.4 mM GTP, retaining a certain degree of curvature and behaving as a nematic phase largely filling the membrane space (**Fig. 4A**). The control by GTP of FtsZ-mts self-organization was found to correlate to the steady-state protein density on the membrane, which was approximately 8-fold higher at 0.04 and 0.4 mM GTP than at 4 mM nucleotide. Interestingly, in the absence of GTP, FtsZ levels at the membrane were two-fold higher than the ones found at 0.04 and 0.4 mM GTP (**Fig. 4B**).

**FIGURE 4.**
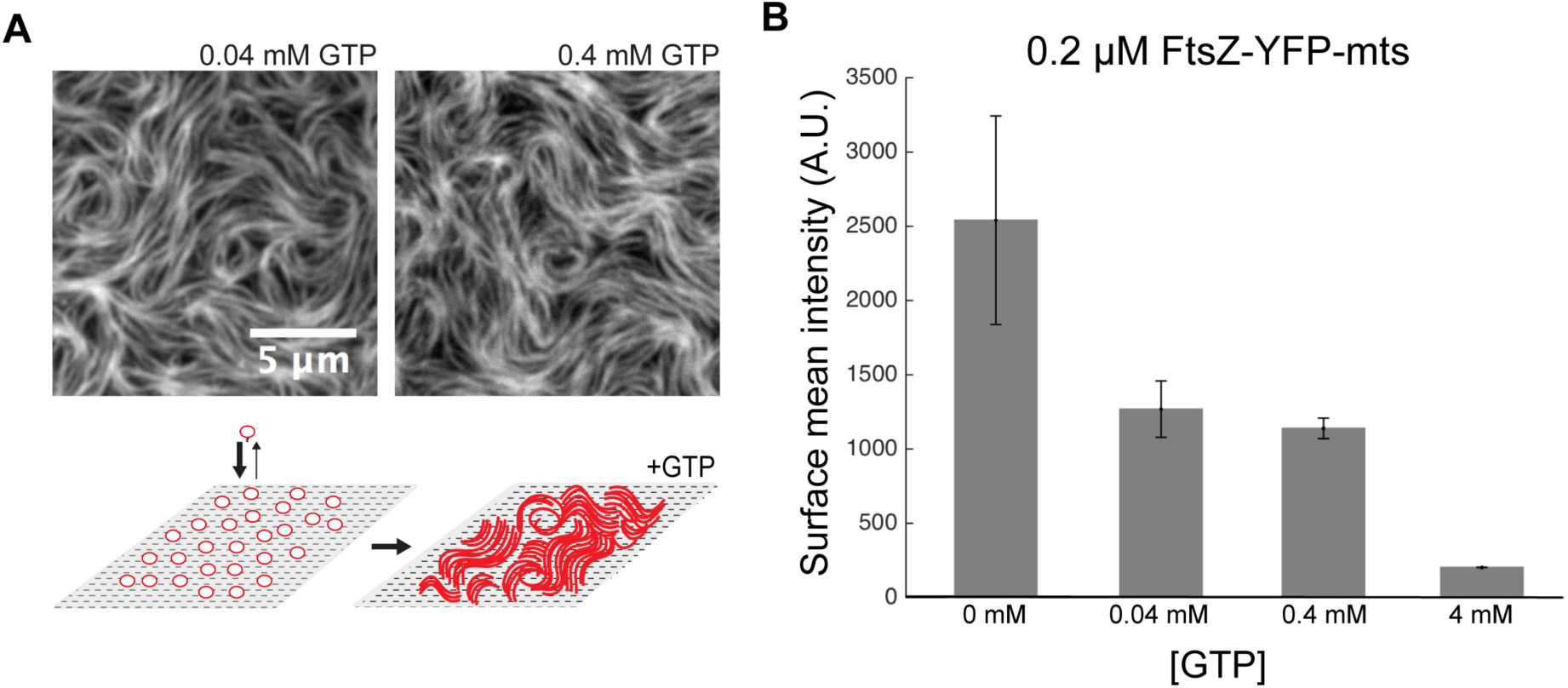
GTP control of FtsZ-mts self-organization is linked to protein density on the membrane. (A) TIRFM images of FtsZ-YFP-mts (0.2 µM) in the presence of 0.04 and 0.4 mM GTP. At GTP concentrations below 1 mM (0.04 mM and 0.4 mM), the available protein seems to rapidly polymerize on the membrane suggesting that in this regime, polymerization on the membrane rules over fast release of protein to solution. Consequently, long filaments with a parallel arrangement are observed at sub-milimolar GTP (schematic representation) corresponding to structures with high protein density. (B) Bar plot of the mean fluorescence intensity for each GTP concentration. Data represents the average of 3 TIRFM imaging experiments. Error bars represent the standard deviation. The mean fluorescence intensity is proportional to the FtsZ-YFP-mts density on the membrane.

The impact of GTP levels on the relative abundance of FtsZ-YFP-mts at the surface was confirmed by single molecule assays (**Fig. 5A**). We determined the probability of protein detachment events at times ≤ t (cumulative probability) for high and low GTP using FtsZ- YFP-mts incubated with fluorescently labelled nanobodies (GFP-Booster-Atto647N, **Fig. 5B**, see Materials and Methods). This revealed that detachment events are faster at high (4mM) than at low (0.04mM) GTP (**Fig. 5C**). These results showed that the residence time of the membrane-targeted FtsZ on the membrane was inversely proportional to the GTP concentration in solution. More importantly, these findings represent compelling evidence that the formation of the dynamic rings at high (4mM) GTP are linked to the faster release of protein from the surface when compared to the slow detachment at lower GTP, when no dynamic circular structures were detected. To rule out the possibility that this effect is due to photobleaching, we also performed the experiments with the illumination intensity reduced by 50%, obtaining the same type of behaviour (**Fig. S5**).

**FIGURE 5.**
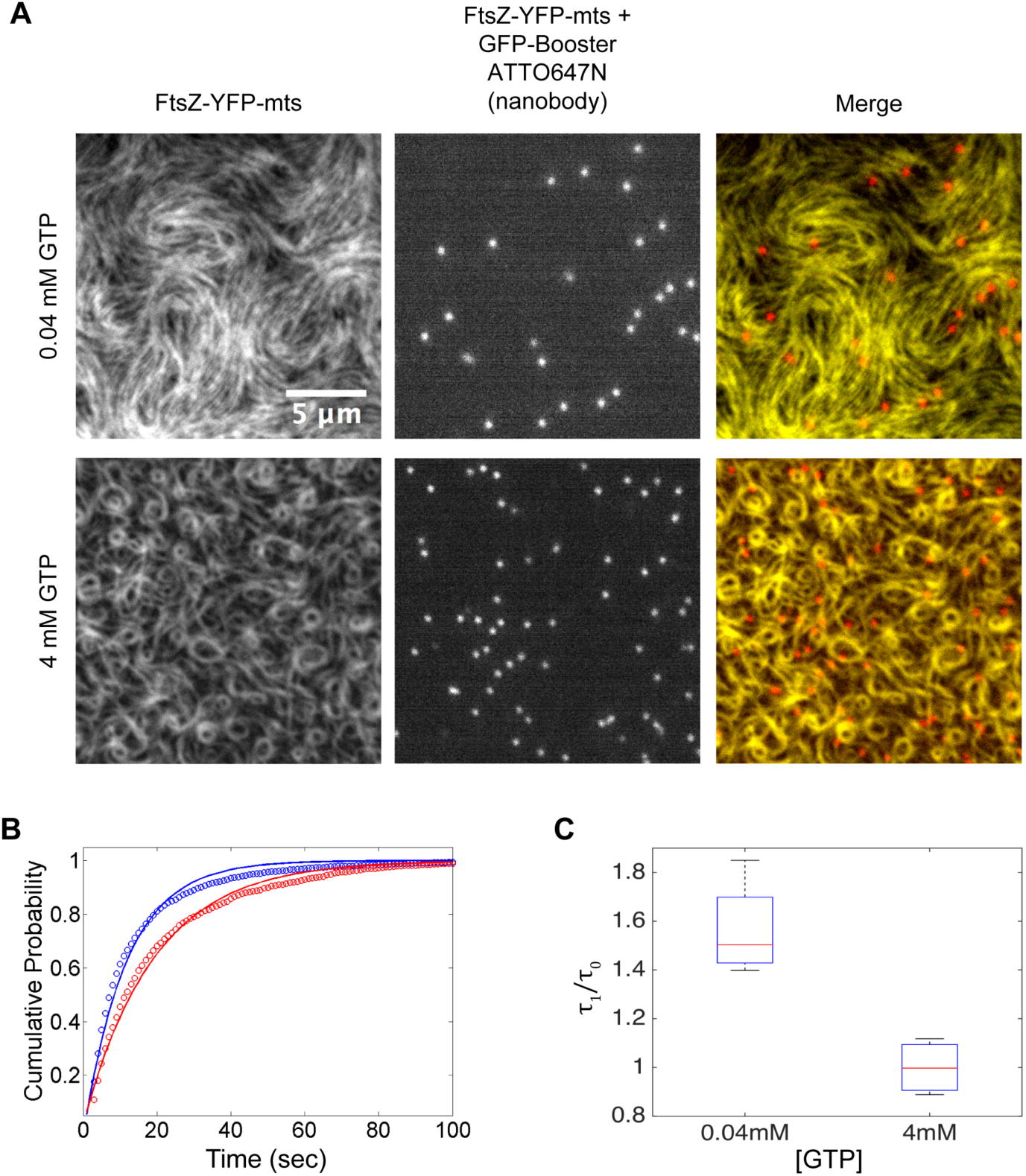
Single molecule dynamics reveals the dependence of the FtsZ-YFP-mts monomer residence time on GTP concentration. Formation of dynamic rings requires frequent turnover of proteins in the membrane-attached filaments. (A) Images of FtsZ-YFP-mts polymers at 0.04 mM and 4 mM GTP doped with 0.8% of FtsZ-YFP-mts incubated with the nanobody GFP-Booster-Atto647N (left panels) are compared with the corresponding images of the nanobodies (mid panels). In the right panels the two images were overlaid (FtsZ-YFP-mts - green, nanobodies - red) (B) Cumulative probability distributions of the events when loss of signal from nanobodies occur. The distributions were measured at 0.04 mM (red circles) and at 4 mM GTP (blue circles) and were fitted to single exponential functions, 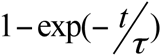, red and blue lines, respectively. The corresponding time scale, τ, represents a generalized residence time that includes both the rate of protein detachment and that of nanobody bleaching (see Materials and Methods). Since all experiments were performed using the same illumination level, the variation of τ with GTP concentration can be used as an indirect measure of the protein detachment rate. While the best fitting curve for the 0.04 mM GTP distribution corresponds to τ ≡τ_1_= 19.2 sec, at 4 mM GTP, τ ≡τ 0 = 11.7 sec, showing that at low GTP concentration FtsZ-YFP-mts is released at a slower rate from the membrane than at high GTP. C) Box plot of τ_1_ / τ_0_ values from 4 independent experiments at each of the two GTP concentrations, showing that the behavior found in (B) is reproduced in different experiments.

## Discussion

In our minimalistic *in vitro* reconstitution study, we found that polymers of an artificially membrane-targeted variant of FtsZ autonomously and without the presence of FtsA self-organize on a supported bilayer upon addition of GTP, to form chiral ring-like dynamic patterns, displaying a clockwise or anti-clockwise protein movement, dependent on whether the membrane attachment was enforced through the C-terminal or N-terminal end of the protein, respectively. The membrane targeting sequence in both cases was taken from MinD, one of the elements of the site-selection MinCDE complex.

Probably the most exciting outcome of this study is the clear dependence of vortex chirality on the positioning of the membrane anchor. It was previously found that liposomes displayed concave or convex deformations when bound to FtsZ-YFP with the mts sequence on the C or N-terminus, respectively (**Fig. 6A**)^13^. To account for their results, it was proposed that the C- or N-terminal attachments were aligned parallel (C) or antiparallel (N) to the direction of curvature of the FtsZ protofilaments, which was, at the same time, perpendicular to the direction of growth, i.e., the filament polarity. To model vortex chirality and its dependence on C/N terminal membrane attachment, we now postulate a new vectorial feature of the FtsZ filaments, namely, that the protein termini are not only located on opposite sides of the filament with respect to growth direction, but also perpendicular to filament curvature (**Fig. 6B**). Assuming that the filament will align on the membrane with the mts anchor facing the bilayer, the chirality of the FtsZ filament dynamics will be clockwise whenever the mts is at the C-terminal and anti-clockwise if located at the N-terminal (**Fig. 6C**). We therefore identify three orthogonal directions for FtsZ filament dynamics on supported membranes: 1) direction of filament growth (polarity), 2) direction of filament curvature, and 3) direction of membrane attachment (**Fig. 6B**).

**FIGURE 6.**
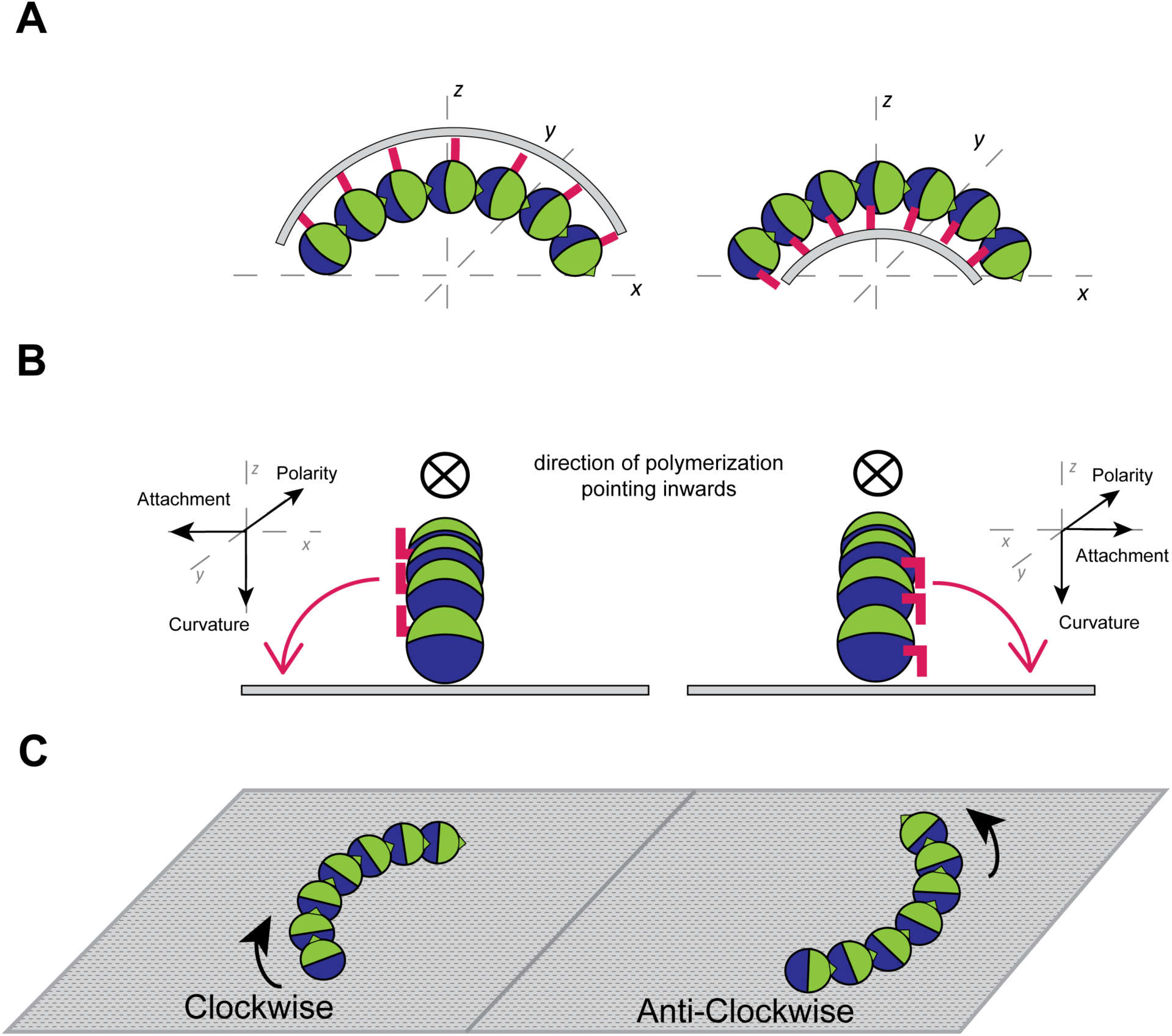
Model for the chirality of FtsZ vortices. A) FtsZ filaments with different terminal mts attached to liposomes^13^. Each protein monomer (spheres) contains a C-domain (green hemisphere) and a N-domain (blue hemisphere) oriented along the backbone of the filament. Each domain ends in a corresponding terminus (C/N, red), that are assumed to be perpendicular to the filament and at 180° from each other. For simplicity, the termini are displayed only when attached to the mts membrane linker. Consistent with the findings of Osawa *et al.* 2009, the filament is curved such that the C-terminals are on the convex side. Moreover, filament assembly occurs preferentially at the C-terminal end. The growth polarity is thus illustrated by the small triangle at the tip of the C-domain. B) Side view of FtsZ filaments vertically aligned on flat supported membranes (an unstable configuration). The main assumption of our model is that the C vs. N terminal attachment induces a symmetry breaking with respect to the direction of polymerization. Accordingly, when the mts linker is at the C-terminal (left panel), as the filament is forced onto a surface, it will prefer to lay down to the left, such that the mts can come into direct contact with the bilayer (red arrow). On the contrary, when the mts linker is at the N-terminal (right panel), the filament will lay down to the right. C) The filaments in B) are now oriented parallel to the surface, such that the mts on either the C-terminal (left panel) or the N-terminal (right panel) is attached to the membrane (in this configuration the terminals are hidden under the filament). Since the active end of the filament is the one that ends with the C-domain, such attachment leads to clockwise or anti-clockwise dynamic chirality, respectively.

The emergence of these reorganizing structures is critically controlled by protein and GTP concentration. The formation of vortices depends on GTP binding and FtsZ polymerization, whereas treadmilling dynamics are driven by destabilization effects, such as fragmentation and annealing, which are strongly linked to GTP hydrolysis. This is clearly observed in the FtsZ variant without GTPase activity that forms static rings compared to the other chimeras with clockwise and counter-clockwise apparent rotation (**Fig. 2**). High GTP enhances the rate of filament turnover and balances FtsZ membrane binding and dissociation, resulting in a steady state destabilization of FtsZ polymer networks on the membrane, and thus driving the self-organization. This suggests that the remodelling of isotropic polymer bundles towards the formation of non-isotropic dynamic patterns of coordinated streams and swingling FtsZ rings is largely governed by the FtsZ surface concentration, which again depends on GTP.

The protein concentration-dependent formation of dynamic FtsZ patterns found in this work nicely correlates with a recent theoretical study showing that protein density at the membrane controls the formation of vortex patterns on membranes in a phase-like behaviour. According to this, there exists a critical regime (“phase”) of protein densities in which curved polar filaments move chirally on a surface and self-assemble into vortex structures. While at low density, filaments travel independently, at high-density regimes they form isotropic networks^20^. Our findings indicate that for different initial adsorption rates, the system reaches the vortex phase at similar concentrations of protein on the membrane (**Fig. 3**). At the corresponding time points at which the same protein surface density is reached, rings have on average the same diameter, treadmilling dynamics and similar thickness. In addition, for sufficiently high concentrations of protein on the bilayer, ring thickening leads to collisions between neighbouring vortices, forcing them to open up, and a new regime ensues consisting of a network of parallel filaments (unpublished data). Interestingly, our results are also compatible with previous atomic force microscopy analysis of the structures formed by FtsZ polymers on mica as a function of protein concentration at the surface^21^. At high protein surface density, FtsZ was found to assemble into a dense layer of polymers that tend to align and aggregate laterally into bundles to fill the entire available surface^21^. Decreasing the surface density led to the formation of curved filaments of smaller radius. The emergence of ring-like structures was more evident at the lowest protein concentration assayed, in which single ring-like structures are the predominant structures^18^.

Thus, our data shows that GTP concentration rules the structural organization and dynamics of FtsZ polymers on the bilayer. While at 4 mM GTP we find the vortex phase, resulting in the formation of dynamic chiral patterns, at lower GTP concentrations (0.4 mM and 0.04 mM) the protein on the membrane forms a relatively static network consisting of parallel filament bundles organized around nodes (**Fig. 4**). The GTP level controls the FtsZ assembly/disassembly cycle linked to GTP hydrolysis, and the rate of filament turnover^22^. Interestingly, the GTPase activity of FtsZ and GTP turnover in solution is higher at GTP > 2 mM^23^. At 4mM GTP (**Fig. 1** and **3**) polymerization and GTP hydrolysis are balanced in a way that filament turnover is overall enhanced, while maintaining the protein concentration on the bilayer in the intermediate range, corresponding to the predicted dynamical vortex phase. Following a rapid imbalance upon GTP addition where most protein monomers in their GTP form detach from the bilayer and form short oligomers in solution, the latter return to the membrane, polymerizing into longer, dynamic filament bundles that extensively exchange proteins with solution via GTP hydrolysis.

Since FtsZ filaments are intrinsically curved, circular structures (and vortices) represent a natural assembly mode at intermediate protein densities^20^. In contrast, at GTP concentrations below 2mM, protein polymerization is the dominant process on the membrane. In this regime, the system evolves directly from the high density of membrane-bound monomers to a crowded, jammed network of mostly parallel filaments (**Fig. 4**).

Previously, the co-reconstitution of FtsZ and FtsA in supported bilayers has revealed the formation of dynamic FtsZ+FtsA vortices that undergo clockwise treadmilling^17^. FtsA was proposed to play a central role in this phenomenon, promoting the anchoring of FtsZ protofilaments to the membrane, and at the same time inducing the disassembly of the membrane-bound FtsZ on the opposite end. As a result, the polymerization-depolymerization process generates the treadmilling and the origin of the apparent rotation. As it appears now, these functionalities of FtsA can be significantly simplified and generalized, as the key requirements for FtsZ self-organization seem to be membrane recruitment in concert with the dynamic, GTP- and surface concentration-regulated turnover of monomers from within the filaments.

The key differences between our system and the one reported by Loose and Mitchinson are twofold. First, while the FtsZ + FtsA swirls prevail for about ten minutes and then disappear by fusing with adjacent travelling streams of bundles, in our experiments the individual FtsZ-YFP-mts rings are persistent and their thickness increases with the concentration of the protein on the bilayer. Second, the apparent rotation around the ring structures is about threefold faster in the FtsZ + FtsA system with an average velocity of 6.5 µm/min^17^, compared to the FtsZ-YFP-mts case where the average velocity is only 2.5 µm/min. Following our quantitative analysis, the fact that Loose and Mitchinson failed to see vortex dynamics in reconstituted FtsZ-YFP-mts without FtsA can be attributed to a relatively high protein concentration of 1.5 µM at which the dynamics also cease in our experiments.

On surfaces, dynamic protein networks that are organized around chiral swirls have been previously observed in both actin-myosin and microtubule-dynein systems^24^. Together with the FtsA+FtsZ^17^, FtsZ-YFP-mts (this work) and the FtsZ on mica systems^21,25^, these form a family of apparently related behaviours despite many microscopic differences. These systems share a set of basic components: a) exchange of proteins between bulk and surface, b) formation of curved filaments on the surface and c) a mechanism that allows the filaments to treadmill on the surface. It is possible that for all such systems the concentration of filaments on the surface is the governing parameter and that a dynamic chiral ring network will be the predominant pattern in the intermediate concentration range. However, while for actin filaments and microtubules, vortex dynamics are due to the corresponding molecular motors, in the FtsZ systems it is directly fuelled by the energy resulting from GTP hydrolysis. Since FtsZ-YFP-mts rings were also shown to exert constricting force on tubular liposomes, FtsZ may be thought of as a primitive form of protein-motor system where it concurrently plays both roles.

Our findings together with the in vitro studies by Loose^17^ and the very recent studies showing the linkage between treadmilling of FtsZ polymers and peptidoglycan synthesis in *E. coli*^26^ and *B. subtilis*^27^ cells, shed new light on the fundamental nature, but also hint to a physiological role of FtsZ treadmilling in bacterial division events. The minimal system we used unambiguously shows that the observed chiral vortices are the result of intrinsic GTP-linked FtsZ polymerization dynamics on the membrane without the need of additional complex interactions with FtsA and ATP. The reduced number of components allowed us to determine the influence of key factors, e.g. the surface density of FtsZ, on the self-organization behavior, thus contributing to the mechanistic understanding of FtsZ treadmilling.

## Materials and Methods

### Mutants

Mutations in *ftsZ-YFP-mts* were constructed using site-directed mutagenesis. FW oligonucleotide was designed using the NEBaseChanger – Substitution (http://nebasechanger.neb.com/). RV oligonucleotide was obtained from the reverser complement sequence of the FW oligonucleotide. 5’- GGTGGTGGTGCCGGTACAGGT-3’ and 5’-ACCTGTACCGGCACCACCACC-3’ oligonucleotides were used to generate FtsZ-YFP-mts-T108A mutant by replacing a Thr in position 108 by an Ala. Briefly, *ftsZ-YFP-mts* was first amplified using the FW and RV oligonucleotides in different PCR reactions, testing three different temperatures: 54ºC, 58.5ºC and 65ºC. In a second PCR reaction, the PCR products from the FW and RV oligonucleotides were mixed; also three different temperatures were tested, 54ºC, 58.5ºC and 65ºC. After digestion with DpnI, the three PCR products were used to transform CH3-Blue competent cells. 5 colonies were picked for DNA extraction and selected for sequencing.

### Protein purification

The FtsZ-YFP-mts, FtsZ*[T108A]-YFP-mts and mts-H-FtsZ-YFP were purified as previously described^12^. Briefly, the protein was expressed from a pET-11b expression vector and transformed into *E. coli* strain BL21. ON over-expression was performed at 20ºC for the proteins FtsZ-YFP-mts and FtsZ*[T108A]-YFP-mts; and at 37ºC for protein mts-H-FtsZ-YFP. Cells were lysed by sonication and separated by centrifugation. Then, protein was precipitated from the supernatant, adding 30% ammonium sulphate and incubating the mixture for 20 min on ice (slow shaking). After centrifugation and resuspension of the pellet, the protein was purified by anion exchange chromatography using a 5x 5ml Hi-Trap Q-Sepharose column (GE Healthcare, 17515601). Purity of the protein was confirmed by SDS-PAGE and mass spectrometry.

### Small unilamellar vesicles (SUVs)

*E. coli* polar lipid extract (Avanti, AL, USA), initially dissolved in chloroform, was desiccated under a gas nitrogen stream. Chloroform traces were further removed by 1-hour vacuum. This lipid film was hydrated with SLB-buffer (50mM Tris-HCl, 150 mM KCl, pH 7.5) to reach a lipid concentration of 4mg ml^−1^. After 10 min sonication, small unilamellar vesicles (SUVs) were obtained.

### Supported lipid bilayers (SLBs)

1.5 glass coverslips (Menzel, Germany) were cleaned in air plasma. Then a plastic ring was attached on a cleaned glass coverslip with a volume of 200 µl using ultraviolet-curable glue (Norland Optical Adhesive 63). SLBs were obtained by the method of vesicle fusion, from SUVs on a glass surface as described elsewhere. The SUV dispersion was diluted in SLB buffer (50  mM Tris-HCl at pH  7.5, 150  mM KCl) to 0.5  mg  ml^−1^, of which 75  µl was added to the reaction chamber. Adding CaCl_2_ to a final concentration of 3  mM promoted vesicle fusion and the formation of a lipid bilayer on the glass. The samples were incubated at 38⁰C for 20 min and then washed with pre-warmed SLB buffer to remove non-fused vesicles. Lastly, 5mM MgCl_2_ are added to the sample.

### Self-organization assays

FtsZ-YFP-mts or FtsZ-YFP-mts-mutants were added to the reaction buffer (50  mM Tris-HCl at pH  7.5, 150  mM KCl and 5  mM MgCl_2_) above the supported lipid membrane in the chamber. The final volume of the samples was ~200  µl. FtsZ-YFP-mts was added to a final concentration of 0.5 µM or 0.2 µM. Polymerization was induced by adding GTP to a final concentration between 0.04 and 4 mM, as indicated in the text.

### Total internal reflection fluorescence microscopy imaging

All experiments were performed on a WF1 GE DeltaVision Elite^®^ total internal reflection fluorescence microscope (GE Healthcare Life Sciences, Germany) equipped with an OLYMPUS 100x TIRF objective (NA 1.49). The UltimateFocus feature of DeltaVision Elite maintains the focus plane constant in time. FtsZ-YFP-mts was excited with a 488 nm diode laser (10 mW, before objective) Fluorescence imaging is performed using a standard FITC filter set. Images were acquired with a PCO sCMOS 5.5 camera (PCO, Germany) controlled by the softWoRx^®^ Software (GE Healthcare Life Sciences, Germany). For time-lapse experiments, images were acquired every 3 or 10 sec, with a 0.05 sec exposure time, with light illumination shuttered between acquisitions.

### Image analysis and processing

Image analysis was carried out in MATLAB 2015 (MATLAB and Image Processing and Computer Vision Toolbox Release 2015a, The MathWorks, Inc., Natick, Massachusetts, United States.) and processing with Fiji/ImageJ (Rasband, W. S., ImageJ, US National Institutes of Health, Bethesda, http://rsb.info.nih.gov/ij/, 1997–2007). Images correspond to average of 5-10 frames from a time-series experiment. For the kymograph analysis, time-series acquisitions were filtered using a standard mean filter and were drift corrected (Image J). A Matlab script allows the user to define a ring by providing two coordinates. Every ring is automatically fitted to a circle with radius r. Then, three trajectories corresponding to three concentric circles having radii r, r+1 and r-1 pixels are determined. At this point, the script will read the time-series data and calculate a kymograph for each time point and trajectory. The final kymograph corresponds to the average of the three different trajectories.

### Single-molecule imaging and residence time measurement

FtsZ-YFP-mts was doped with 0.8% FtsZ-YFP-mts previously incubated 1:1 with the nanobody GFP-Booster Atto647N (ChromoTek, Germany) for at least 1 hour at 4°C under agitation. FtsZ-YFP-mts + GFP-Booster Atto647N was excited with a 640nm diode laser, (50mW and 100mW, before objective). Single molecule imaging was performed using a standard Cy5 filter set. After 10-15 minutes upon GTP addition, a significant number of spots in the single molecule channel (Atto647N) were observed and imaged at a rate of 1 image per second with 0.5 sec exposure time. To determine the position of every single molecule and calculate its residence time, we employed a Matlab routine designed by Weimann & Ganzinger ^28^. Briefly, a band pass filter was used to remove low and high frequency noise. Then, single molecule positions with intensity above a user-defined threshold were determined by their brightness-weighted centroid. The detection algorithm is highly efficient for detecting particles with a signal to noise ratio above 3. The user-defined threshold was chosen to detect the largest number of spots and kept constant for all experiments. Single molecules were tracked among consecutive frames in an area given by a radius of 10 pixels (pixel size = 0.042µm). Thus, the residence time is defined as the time that the particle stays in this area before its signal vanishes.

There are two events that may lead to the loss of the signal from an individual nanobody: (1) detachment from the membrane, or (2) bleaching of the dye. Although our experiment cannot distinguish between these two events, we measured the effect of GTP concentration on the rate of signal loss maintaining the same illumination conditions. To this end, we calculated the probability as a function of time *t* to obtain a loss of signal event at times ≤ t (cumulative probability, **Fig. 5B**). We found that a single exponential function, 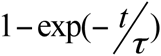, provides a good fit to the measured cumulative probabilities with a corresponding time scale τ, that represents a generalized residence time. Since the bleaching rate was the same in all experiments (same illumination conditions), the variation in τ is due to the effect of GTP concentration on the time that FtsZ-YFP-mts remains attached to the membrane (**Fig. 5C**).

The cumulative probability was measured for 4 different experiments at high GTP concentration, 4 mM, with N = 3519, 7112, 2639, 500 (N = the total number of events in each experiment) and another 4 at low GTP concentration, 0.04 mM, with N = 1331, 1630, 463, 1704. Residence times shorter than 2 seconds are below the accuracy of our method and were not included in the statistics. To rule out possible artefacts due to photo-bleaching, we repeated our measurements using only half the previous laser intensity and obtained a similar ratio between the residence times at the two GTP concentrations as in **Fig. 5C** (**Fig. S5**).

## Contributions

D.R., D.A.G.-S. and A.R. performed research; all authors designed research, analyzed data and wrote the paper.

## Acknowledgements

We are grateful to M. Osawa and H. Erickson for the mts-H-FtsZ-YFP and FtsZ-YFP-mts plasmids. We thank I. Fishov and the members of G.R. lab for discussions and support; M. Schaper and K. Andersson for molecular cloning and purification assistance; Dr. K. Ganzinger for providing code and advice in the single molecule experiments; P. Blumhardt for suggesting the use of the nanobody; N. Ropero for assistance with protein purification; C. Alfonso and J.R. Luque for analytical ultracentrifugation analysis; D.R. and D.A.G.-S are supported by a DFG Fellowship through the Graduate School of Quantitative Biosciences Munich (QBM), and A.R. is funded though the German-Israeli Foundation for Scientific Research and Development.

This work was supported by Human Frontiers Science Program Grant RGP0050/2010 (to G.R. and P.S.); the German-Israeli Foundation for Scientific Research and Development Grant No. 1160-137.14/2011 (to M.F. and P.S.); Gottfried Wilhelm Leibniz-Program of the DFG and the MaxSynBio consortium, which is jointly funded by the Federal Ministry of Education and Research of Germany and the Max Planck Society (to P.S); Spanish Government Grant BFU2014-52070-C2-2-P (to G.R.); and Israel Academy of Science and Humanities Grant No. 1701/13 (to M.F.).

